# The flower flies and the unknown diversity of Drosophilidae (Diptera): a biodiversity inventory in the Brazilian fauna

**DOI:** 10.1101/402834

**Authors:** Hermes J. Schmitz, Vera L. S. Valente

## Abstract

Diptera is a megadiverse order, reaching its peak of diversity in Neotropics, although our knowledge of dipteran fauna of this region is grossly deficient. This applies even for the most studied families, as Drosophilidae. Despite its position of evidence, most aspects of the biology of these insects are still poorly understood, especially those linked to natural communities. Field studies on drosophilids are highly biased to fruit-breeders species. Flower-breeding drosophilids, however, are worldwide distributed, especially in tropical regions, although being mostly neglected. The present paper shows results of a biodiversity inventory of flower-breeding drosophilids carried out in several localities in Brazil, based on samples of 125 plant species, from 47 families. Drosophilids were found in flowers of 56 plant species, from 18 families. The fauna discovered showed to be highly unknown, comprising 28 species, 12 of them (>40%) still undescribed. Not taking in account opportunist species, two thirds of the diversity exclusive from flowers were undescribed. The *Drosophila bromeliae* species group was the most representative taxon, with eight species (six undescribed), including four polyphagous and four *Solanum*-specialised species. This specialisation on *Solanum* is reported for the first time for *Drosophila*. Other taxa of restrict flower-breeding drosophilids were the *Drosophila lutzii* species group and the genus *Zygothrica*. Some specimens of the genera *Cladochaeta, Rhinoleucophenga* and *Scaptomyza* were found, but their relations to flowers are unclear. Additionally, ten species of ample niche was found using flowers opportunistically. Localities and host plants are recorded for all species collected.

## INTRODUCTION

Dealing with the huge diversity of living forms is one of the major challenges for biological sciences. Estimates range from three to 100 millions species, 5-10 millions being the more plausible ones, and a great portion of this biodiversity is experiencing high levels of human-caused threats (Myers *et al.* 2000; May 2010). The invertebrate, and especially, insect diversity account for a great proportion of the total biodiversity. Almost one million species of insects are formally described worldwide and Brazil, with a huge territory and an outstanding environmental heterogeneity, is a key country in this context, housing the highest insect diversity of the world. Estimates indicate that Brazilian fauna harbours 500 thousands to one million insect species, around 10% of the insect diversity of the Earth (Rafael *et al.* 2009).

Diptera is one of the megadiverse orders, comprising alone about 10% of the world’s biodiversity. Neotropics appear to be the most diverse region in these insects, although our knowledge of Neotropical dipteran fauna remains grossly deficient (Brown 2005). This is true even for the most studied families, as Drosophilidae. For more than one century, species of *Drosophila* Meigen, 1830 are largely used as model organisms for a variety of studies, especially in genetical and evolutionary research (Powell 1997). This undoubtedly has brought an exceptional attention for this group of organisms, stimulating early studies on taxonomy and natural history of the family (Sturtevant 1916, 1921; Duda 1925, 1927). In spite of this position of evidence, most aspects of the biology of these insects are poorly understood, especially those linked to natural communities.

The diversity of the family is also barely assessed. This family is one of the larger in Diptera, possessing more than 4,000 described species (Yassin 2013), besides a great number waiting for recognition. The Brazilian fauna of drosophilids has for long being researched (Dobzhansky & Pavan 1943, 1950; Pavan & Cunha 1947) and more than three hundred species are recorded (Gottschalk *et al.* 2008). Therefore, records of drosophilids in Brazilian territory are strikingly concentrated to some regions, and inventories for most regions are still lacking (Chaves & Tidon 2008; Gottschalk *et al.* 2008). Furthermore, samples are greatly biased to fruit-breeders and species attracted to banana-baited traps.

Brncic (1983) listed 140 species of drosophilids associated to flowers in some manner worldwide. This number includes from very generalist species, which use flowers opportunistically, to specialised species, which depend on the flowers in all stages of life cycle. These last species are rarely detected in the traditional samples taken with fruit baits. In Neotropics, the best studied flower-specialised species belong to the *Drosophila flavopilosa* species group (Brncic 1962a, 1966, 1978; Santos & Vilela 2005; Robe *et al*. 2013), restrict to flowers of plants of the genus *Cestrum* (Solanaceae). Other notable *Drosophila* anthophilic taxa in Neotropical Region are the *D. bromeliae* and *D. lutzii* species groups, both widespread in Neotropics and associated to flowers of several botanical families (Chassagnard & Tsacas 1992; Silva & Martins 2004; Grimaldi 2016; Vilela & Prieto 2018); the *D. onychophora* and *D. xanthopallescens* species groups, reported respectively to northern Andes and Central America (Pipkin 1964; Hunter 1979), and some species of uncertain affinities associated to Araceae inflorescences (Tsacas & Chassagnard 1992; Vaz *et al.* 2014, 2018; Llangarí & Rafael 2017). Reports from other genera are also known, especially for *Zygothrica* (Grimaldi 1987; Endara *et al.* 2010; Fonseca *et al.* 2017) and the poorly known *Laccodrosophila* and *Zapriothrica* (Heed *et al.* 1960).

Anthophilic drosophilids, therefore, are present worldwide, especially in tropical regions. Other noteworthy examples are the *Scaptodrosophila aterrima* species group in Africa (Tsacas *et al.* 1988), the subgenus *Exalloscaptomyza* Hardy, 1965 of *Scaptomyza* Hardy, 1949, in Hawaii (Montague & Kaneshiro 1982; Starmer & Bowles 1994), *Scaptodrosophila hibisci* (Bock, 1977) and *S. aclinata* (McEvey & Barker, 2001), in Australia (Cook *et al.* 1977; McEvey & Barker 2001), the *Drosophila elegans* species subgroup (*melanogaster* group) (Sultana *et al.* 1999; Suwito *et al.* 2002), the genus *Colocasiomyia* de Meijere, 1924 (Carson & Okada 1980; Sultana *et al.* 2006; Fartyal *et al.* 2013) and the genus *Arengomyia* Yafuso and Toda, 2008 (Yafuso *et al.* 2008), in Asia.

In addition to being normally neglected in diversity inventories, the nature of the interaction between drosophilids and flowers is poorly known in most cases, since flies may show different levels of dependence on flowers in all or just part of their lyfe-cycles. In an attempt to both decrease the inventory gap and comprehend better such interactions, in the present paper we deal exclusively with flower-breeding species, *i. e.*, those that use flowers as oviposition sites. So, we report here a diversity inventory of the Brazilian flower-breeding drosophilid fauna, based on an ample but not exhaustive sampling effort on a variety of plants, providing information on host plants, localities, including many new records, and the discovery of a highly unknown diversity, comprising mainly undescribed species.

## MATERIAL AND METHODS

The present paper comprises samples taken in several localities throughout the Brazilian territory (Figure 1) between the years of 2006 and 2010. Most of the collections were carried out in the states of Rio Grande do Sul and Santa Catarina, the two southernmost Brazilian states. Additional samples were taken from the states of Paraná, São Paulo, Minas Gerais, Bahia, Pernambuco and Pará. Coordinates were taken with GPS and are shown in Results.

**Figure 1:**
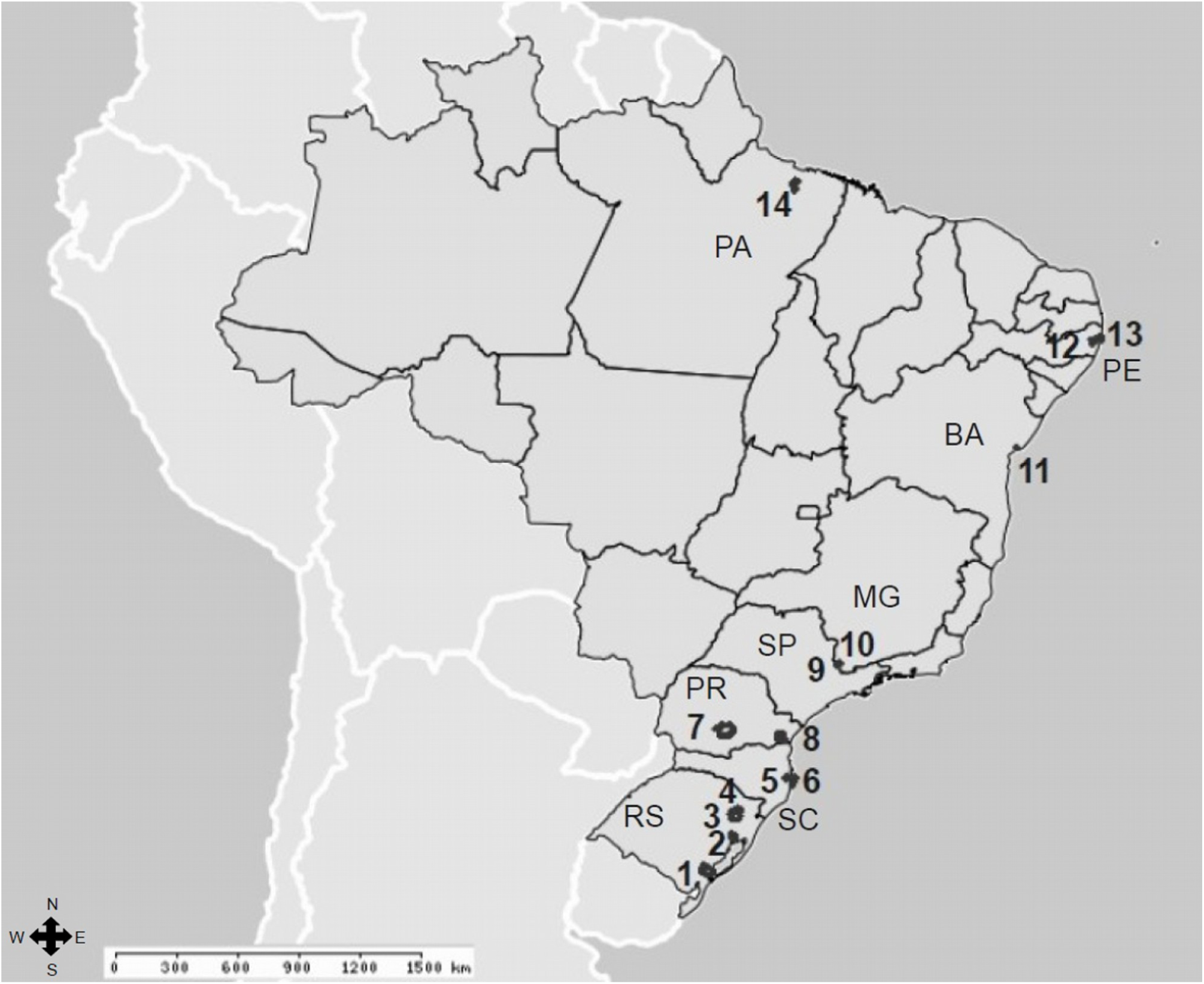
Map of Brazil showing the states and municipalities where samples of this study were taken: RS – Rio Grande do Sul: 1. Pelotas, 2. Porto Alegre, 3. Caxias do Sul, 4. Flores da Cunha; SC – Santa Catarina: 5. Antônio Carlos, 6. Florianópolis; PR – Paraná: 7. Guarapuava, 8. São José dos Pinhais; SP – São Paulo: 9. Águas de Lindóia; MG – Minas Gerais: 10. Monte Sião; BA – Bahia: 11. Itaparica; PE – Pernambuco: 12. Vitória de Santo Antão, 13. Recife; PA – Pará: 14. Belém.

Flowers of 125 species of plants, from 47 families, were collected. They were detached during anthesis directly from the plants, or collected as recently fallen flowers in the ground. They were put in plastic bags and taken to laboratory, where they were transferred to vials with vermiculite, closed with foam stoppers and kept at 25°C, being inspected for the emergence of drosophilid imagines. In this case, the specimens were aspirated, aged for few days in standard culture medium and identified by external morphology and terminalia, consulting specialised literature (Val 1982; Vilela 1984, 1986; Grimaldi 1987, 2016; and others). For analysis of terminalia, postabdomens were prepared following Bächli *et al.* (2004) and stored in microvials with glycerol. Since many species found are probably undescribed and are currently being object of further studies, voucher specimens will be pinned (double-mounted), associated with their respective terminalia in glycerol and deposited later, after taxonomic study.

Plant species were identified after Souza and Lorenzi (2005), references therein and other specialised literature. Eventually, some plants were identified by specialists (see Acknowledgments). Plants scientific names follow APG IV (2016) as in Plants of the World Online website (plantsoftheworldonline.org) or Flora do Brasil 2020 (floradobrasil.jbrj.gov.br).

## RESULTS

### Fauna and flora

From the 125 species of plants sampled, 56 proved to be host for flower-breeding drosophilids, spread in 18 botanical families. Plant species with no emergence of drosophilids in our collections are listed in Appendix S1 in Supporting Information.

All over, 28 species of drosophilids were found. The species found were divided, *grosso modo*, in three categories: (1) “restrict” - species which were caught repeatedly and seem to use living flowers as an exclusive or at least constant breeding site (Table 1); (2) “uncertain biology” - species of genera with poorly known biology and collected only occasionally (Table 2); and (3) “opportunists” - species known to be mainly fruit-breeders, using flowers only incidentally or as a secondary resource, and preferentially fallen flowers (Table 3).

**Table 1:**
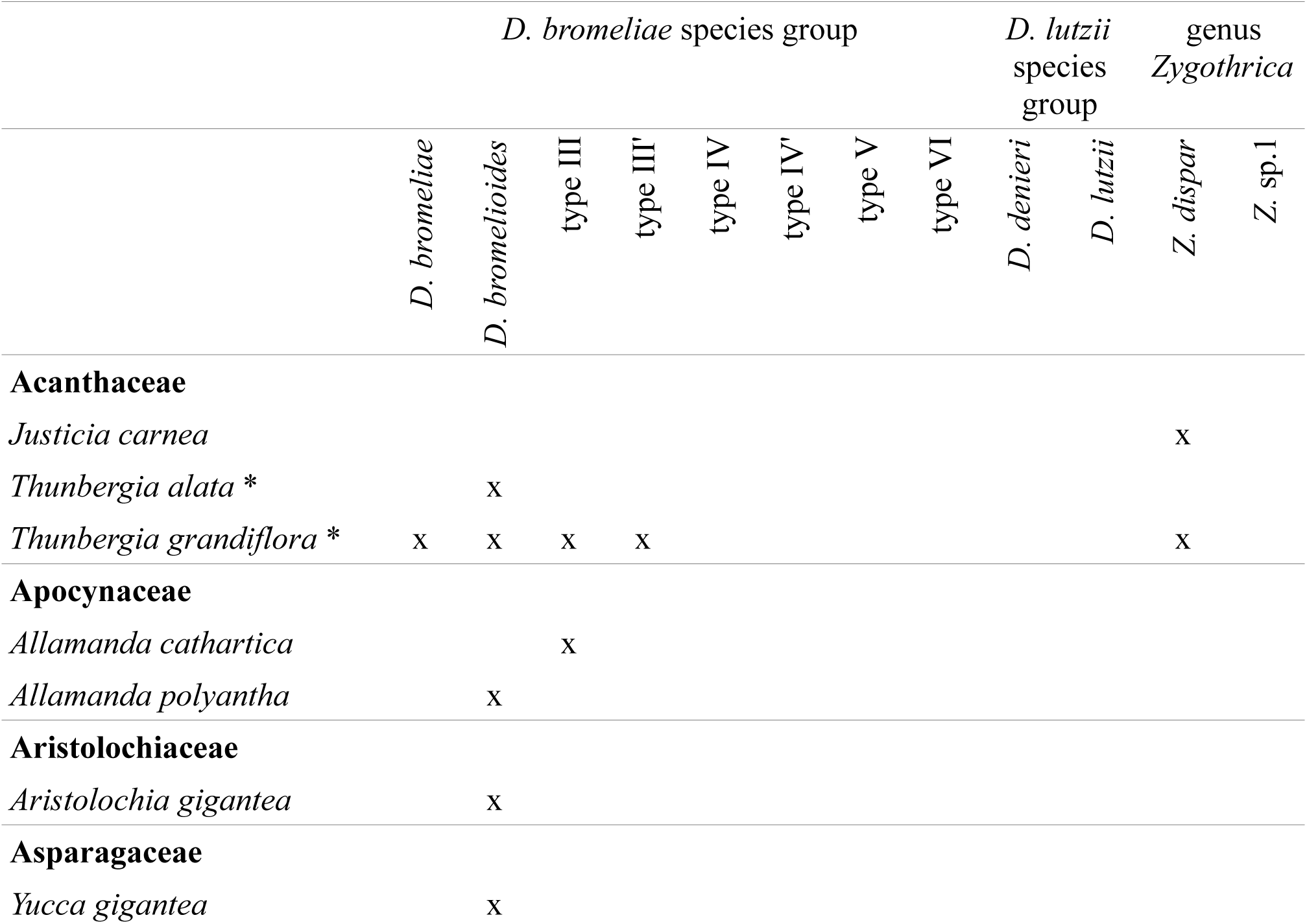
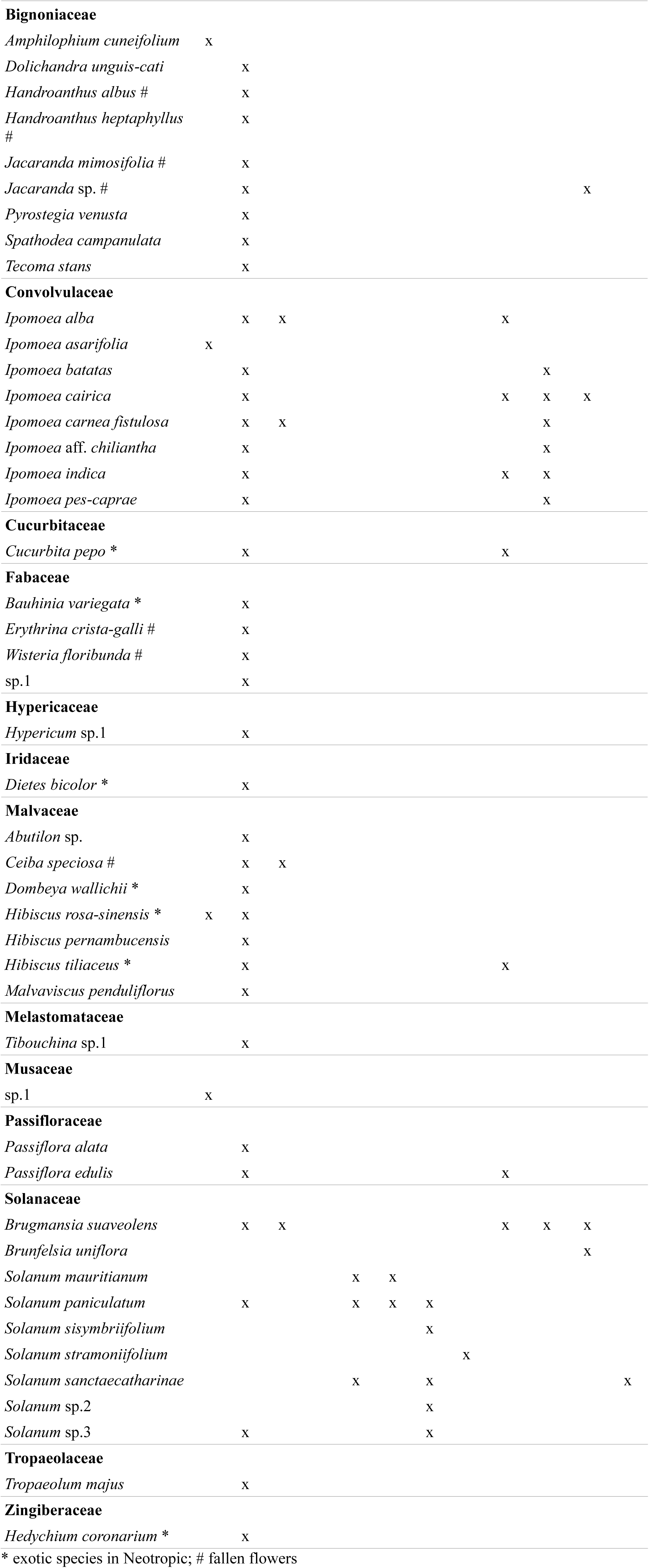
Host plants for Brazilian restrict flower-breeding species of drosophilids found in the present survey.

**Table 2:** Host plants for Brazilian species of drosophilids of uncertain biology found in flowers in the present survey.

|  | *C. bupeo* | *C. sp.1* | *R. joaquina* | *S. sp.1* | *S. sp.2* | *S. sp.3* |
| --- | --- | --- | --- | --- | --- | --- |
| **Bignoniaceae** |  |  |  |  |  |  |
| *Handroanthus albus* # |  |  |  | x |  |  |
| *Handroanthus heptaphyllus* # |  |  |  | x | x | x |
| *Pyrostegia venusta* | x |  |  | x |  |  |
| **Bromeliaceae** |  |  |  |  |  |  |
| *Dyckia encholirioides* |  |  | x |  |  |  |
| **Malvaceae** |  |  |  |  |  |  |
| *Dombeya wallichii* * |  |  |  | x |  |  |
| *Dombeya tiliacea* * |  | x |  |  |  |  |

\* exotic species in Neotropic; # fallen flowers

**Table 3:** Host plants for Brazilian opportunist species of drosophilids found in flowers in the present survey.

|  | *D. cardinoides* | *D. griseolineata* | *D. mediotriata* | *D. mediovittata* | *D. melanogaster* * | *D. paulistorum* | *D. simulans* * | *D. sticta* | *D. tripunctata* gr. sp.1 | *Z. indianus* * |
| --- | --- | --- | --- | --- | --- | --- | --- | --- | --- | --- |
| **Acanthaceae** |  |  |  |  |  |  |  |  |  |  |
| *Thunbergia grandiflora* * |  |  |  |  | x |  |  |  |  |  |
| **Apocynaceae** |  |  |  |  |  |  |  |  |  |  |
| *Allamanda polyantha* |  |  |  |  |  | x |  |  |  |  |
| **Aristolochiaceae** |  |  |  |  |  |  |  |  |  |  |
| *Aristolochia gigantea* |  |  |  |  | x |  |  |  |  |  |
| **Asparagaceae** |  |  |  |  |  |  |  |  |  |  |
| *Yucca gigantea* # |  |  |  |  |  |  | x |  |  | x |
| **Bignoniaceae** |  |  |  |  |  |  |  |  |  |  |
| *Handroanthus albus* # |  | x |  |  |  |  |  | x |  |  |
| *Handroanthus heptaphyllus* # |  |  |  | x |  |  | x |  |  |  |
| **Fabaceae** |  |  |  |  |  |  |  |  |  |  |
| *Erythrina speciosa* # |  |  |  |  |  |  |  |  | x |  |
| **Malvaceae** |  |  |  |  |  |  |  |  |  |  |
| *Ceiba speciosa* # | x |  |  |  |  |  |  | x |  |  |
| *Hibiscus pernambucensis* # |  |  | x |  |  |  |  |  |  |  |

\* exotic species in Neotropic; # fallen flowers

### Localities

The localities where which species was found are listed below. They are given from North to South, in the following format: State: Municipality, Locality (latitude; longitude; altitude, when available).

### (1) “Restrict” flower-breeding species

#### *Drosophila bromeliae* species group

##### *D. bromeliae* Sturtevant, 1921

Pará: Belém, Reserva do Mocambo (01°26’34.35”S; 48°24’35.59”W; 40m); Pernambuco: Recife, UFPE campus (08°03’14”S; 34°57’00”W); Recife, UFRPE campus (08°00’45”S; 34°56’57”W); Vitória de Santo Antão, IFPE/Campus Vitória (08°06’04.1”S; 35°17’41.3”W; 152m); Mato Grosso: Tangará da Serra, “Park in town center” (14°04’38”S; 57°03’45”W) (specimens collected [“emerged from flowers from Convolvulaceae”] and cited as *Drosophila* sp.1 by Blauth & Gottschalk 2007 and reviewed by HJS).

##### *D. bromelioides* Pavan and Cunha, 1947

São Paulo: Águas de Lindóia, Morro do Cruzeiro (22°28’43”S; 46°37’34”W); Paraná: Guarapuava, Parque do Lago (25°24’9.099”S; 51°28’30.82”W); Santa Catarina: Governador Celso Ramos, Anhatomirim Island (27°25’38.09”S; 48°33’53.89”W); Florianópolis, Ratones Grande Island (27°28’20.20”S; 48°33’43.00”W); Antônio Carlos, Centro (27°31’03.373”S; 48°46’18.23”W); Florianópolis, Ponta do Coral (27°34’23”S; 48°31’41”W); Florianópolis, Santa Mônica (27°35’34”; 48°30’53”W); Florianópolis, Morro da Lagoa da Conceição (27°35’40.9”S; 48°28’41.6”); Florianópolis, UFSC campus (27°36’08.127”S; 48°31’30.39”W); Florianópolis, Joaquina (27°36’43.9”S; 48°26’36.3”W); Rio Grande do Sul: Caxias do Sul, Grupo de Escoteiros Saint Hilaire (29°09’38.2”S; 51°08’31.7”W; 706m); Caxias do Sul, Marcopolo/BR-116 road (29°11’06.4”S; 51°10’23.8”W; 695m); Porto Alegre, Parque Farroupilha (30°02’13.0”S; 51°13’06.0”W); Porto Alegre, Parque Maurício Sirotsky Sobrinho (30°02’34”; 51°13’55”W); Porto Alegre, UFRGS/Campus do Vale (30°04’12.4”S; 51°07’06.9”W); Porto Alegre, Parque Gabriel Knijnick (30°06’12”S; 51°12’08”W); Pelotas, Balneário dos Prazeres (31°43’28.4”S; 52°11’59.5”W; 15m, and 31°44’06.5”S; 52°12’54.1”W; 21m); Pelotas, near Laranjal (31°45’28.4”S; 52°15’42.8”W; 15m); Pelotas, Fragata (31°45’18.3”; 52°23’01.1”W; 62m); Pelotas, Fragata, near Arroio Fragata (31°45’29.5”S; 52°24’05.05”W; 62m).

#### Type III

Santa Catarina: Antônio Carlos, Rachadel (27°29’52.8”S; 48°47’38.6”W); Rio Grande do Sul: Porto Alegre, Parque Marinha do Brasil (30°03’16.3”S; 51°13’57.8”W); Porto Alegre, UFRGS/ Campus do Vale (30°04’12.4”S; 51°07’06.9”W); Pelotas, Balneário dos Prazeres (31°43’28.4”S; 52°11’59.5”W; 15m).

#### Type III’

Pernambuco: Recife, UFPE campus (08°03’14”S; 34°57’00”W).

#### Type IV

Santa Catarina: Florianópolis, Córrego Grande (27°35’37.6”S; 48°29’37.0”W); Rio Grande do Sul: Cruz Alta, CEPPA (28°34’11”S; 53°36’53”W) (*D.* type IV of Hochmüller *et al.*, 2009); Flores da Cunha, RS-122 road (29°01’27.9”S; 51°11’41.0”W; 729m, and 29°01’37.4”S; 51°11’43.7”W; 733m); Porto Alegre, UFRGS/Campus do Vale (30°04’12.4”S; 51°07’06.9”W).

#### Type IV’

Bahia: Itaparica, Itaparica Island, (12°54’18”S; 38°40’01”W); Minas Gerais: Monte Sião, Centro (22°25’36.16”S; 46°34’10.43”W); São Paulo: Águas de Lindóia, Morro do Cruzeiro (22°28’43”S; 46°37’34”W); Paraná: São José dos Pinhais, Águas Belas (25°32’09.71”S; 49°10’52.87”W); Santa Catarina: Antônio Carlos, Rachadel (27°29’48.02”S; 48°47’39.70”W).

#### Type V

Santa Catarina: Florianópolis, Córrego Grande (27°35’37.6”S; 48°29’37.0”W); Rio Grande do Sul: Porto Alegre, UFRGS/Campus do Vale (30°04’12.4”S; 51°07’06.9”W); Pelotas, Balneário dos Prazeres (31°43’28.4”S; 52°11’59.5”W; 15m, and 31°44’06.5”S; 52°12’54.1”W; 21m).

#### Type VI

Pará: Belém, Reserva do Mocambo (01°26’34.35”S; 48°24’35.59”W; 40m).

#### ***Drosophila lutzii* species group (=subgenus *Phloridosa* of *Drosophila*)**

##### *D. denieri* Blanchard, 1938

Santa Catarina: Antônio Carlos, Centro (27°31’03.373”S; 48°46’18.23”W); Florianópolis, Ponta do Coral (27°34’23”S; 48°31’41”W); Rio Grande do Sul: Caxias do Sul, Marcopolo/BR-116 road (29°11’01.3”S; 51°10’14.4”W; 701m, and 29°11’06.4”S; 51°10’23.8”W; 695m); Porto Alegre, Parque Maurício Sirotsky Sobrinho (30°02’34”; 51°13’55”W); Porto Alegre, UFRGS/Campus do Vale (30°04’12.4”S; 51°07’06.9”W); Pelotas, Fragata (31°45’18.3”; 52°23’01.1”W; 62m); Pelotas, Fragata, near Arroio Fragata (31°45’29.5”S; 52°24’05.05”W; 62m).

##### *D. lutzii* Sturtevant, 1916

São Paulo: Águas de Lindóia, Centro (22°28’17”S; 46°37’36”W); Águas de Lindóia, Morro do Cruzeiro (22°28’43”S; 46°37’34”W); Paraná: Guarapuava, Parque do Lago (25°24’9.099”S; 51°28’30.82”W); Santa Catarina: Governador Celso Ramos, Anhatomirim Island (27°25’38.09”S; 48°33’53.89”W); Florianópolis, UFSC campus (27°36’08.127”S; 48°31’30.39”W); Rio Grande do Sul, Caxias do Sul, Marcopolo/BR-116 road (29°11’01.3”S; 51°10’14.4”W; 701m); Rio Grande do Sul: Porto Alegre, Parque Maurício Sirotsky Sobrinho (30°02’34”; 51°13’55”W); Porto Alegre, UFRGS/Campus do Vale (30°04’12.4”S; 51°07’06.9”W).

#### Genus *Zygothrica* Wiedemann, 1830

##### *Z. dispar* (Wiedemann, 1830)

Santa Catarina: Florianópolis, Morro da Lagoa da Conceição (27°35’40.9”S; 48°28’41.6”W; 200m); Rio Grande do Sul: Porto Alegre, UFRGS/Campus do Vale (30°04’12.4”S; 51°07’06.9”W).

##### *Z.* sp.1

Rio Grande do Sul: Porto Alegre, UFRGS/Campus do Vale (30°04’12.4”S; 51°07’06.9”W).

### (2) “Uncertain biology”

#### *Cladochaeta bupeo* Grimaldi and Nguyen, 1999

Rio Grande do Sul: Porto Alegre, UFRGS/Campus do Vale (30°04’12.4”S; 51°07’06.9”W).

#### *Cladochaeta* sp.1

(*Cladochaeta nebulosa* species group). Rio Grande do Sul: Porto Alegre, Parque Farroupilha (30°02’13.0”S; 51°13’06.0”W).

#### *Rhinoleucophenga joaquina* Schmitz *et al.*, 2009

Santa Catarina: Florianópolis, Parque Municipal das Dunas da Lagoa da Conceição, Joaquina (27°38’S; 48°28’W).

#### *Scaptomyza* sp.1

(subgenus *Mesoscaptomyza*). Rio Grande do Sul: Porto Alegre, Parque da Marinha (30°03’40.75”S; 51°13’40.75”W); Porto Alegre, UFRGS/Campus do Vale (30°04’12.4”S; 51°07’06.9”W).

#### *Scaptomyza* sp.2

(subgenus *Parascaptomyza*). Rio Grande do Sul: Porto Alegre, UFRGS/Campus do Vale (30°04’12.4”S; 51°07’06.9”W).

#### *Scaptomyza* sp.3

(subgenus *Mesoscaptomyza*). Rio Grande do Sul: Porto Alegre, UFRGS/Campus do Vale (30°04’12.4”S; 51°07’06.9”W).

### (3) “Opportunists”

#### *D. cardinoides* Dobzhansky and Pavan, 1943

Rio Grande do Sul: Porto Alegre, UFRGS/Campus do Vale (30°04’12.4”S; 51°07’06.9”W).

#### *D. griseolineata* Duda, 1927

Rio Grande do Sul: Porto Alegre, UFRGS/Campus do Vale (30°04’12.4”S; 51°07’06.9”W).

#### *D. mediostriata* Duda, 1925

Santa Catarina: Florianópolis, Ponta do Coral (27°34’23”S; 48°31’41”W).

#### *D. mediovittata* Frota-Pessoa, 1954

Rio Grande do Sul: Porto Alegre, UFRGS/Campus do Vale (30°04’12.4”S; 51°07’06.9”W).

#### *D. melanogaster* Meigen, 1830

Pernambuco: Recife, UFPE campus (08°03’14”S; 34°57’00”W); Santa Catarina: Florianópolis, UFSC campus (27°36’08.127”S; 48°31’30.39”W).

#### *D. paulistorum* Dobzhansky and Pavan, 1949

Rio Grande do Sul: Porto Alegre, UFRGS/Campus do Vale (30°04’12.4”S; 51°07’06.9”W).

#### *D. simulans* Sturtevant, 1919

Rio Grande do Sul: Porto Alegre, UFRGS/Campus do Vale (30°04’12.4”S; 51°07’06.9”W).

#### *D. sticta* Wheeler, 1957

Santa Catarina: Antônio Carlos, Centro (27°31’03.373”S; 48°46’18.23”W); Rio Grande do Sul: Porto Alegre, UFRGS/Campus do Vale (30°04’12.4”S; 51°07’06.9”W).

#### *D. tripunctata* group sp.1

Rio Grande do Sul: Porto Alegre, Parque Farroupilha (30°02’13.0”S; 51°13’06.0”W).

#### *Zaprionus indianus* Gupta, 1970

Rio Grande do Sul: Porto Alegre, Parque Farroupilha (30°02’13.0”S; 51°13’06.0”W).

## DISCUSSION

The flower-breeding drosophilid fauna of Brazil proved to be strikingly unknown. From the 28 species found, 12 were species of unknown identity and probably still undescribed at the moment of collection, *i. e.*, ∼ 43%. The ignorance in respect to this fauna is still more pronounced if one takes by side the opportunist fruit-breeders species and considers only the exclusive fauna of the flowers (our “restrict” and “uncertain biology” categories): 12 undescribed species out of 18, *i. e.*, two thirds of them. Undescribed species of drosophilids in Brazilian fauna are not uncommon, even in surveys based on banana-baited traps. Medeiros and Klaczko (2004) estimated that half of the *Drosophila* species of the state of São Paulo, the best sampled Brazilian state, was still undescribed. A high number of undescribed species was also collected in Santa Catarina, other relatively well-studied state, by Gottschalk *et al.* (2007) and Döge *et al.* (2008). The diversity of flower-breeding drosophilids, although clearly lower than that associated to fruits, appears to be even more unknown. This is especially astonishing considering that Drosophilidae is one of the most studied taxa of Diptera. Brown (2005) estimated that for the largest Diptera families, the rate of new described species is only 7.6 per year in the Neotropical region, and more than 1,000 years would be necessary to describe all species of Diptera in the world. Furthermore, a described species does not mean necessarily a well-known species: several described species found in the present survey are recorded for the first time in some Brazilian states or even in South America. In fact, drosophilid records in Brazil are strongly biased towards fruit-feeding species of *Drosophila*, and to southeastern and southern localities (more exactly, near the main universities, and the present survey is no exception); moreover, about one third of the species recorded in Brazil are known for a single locality (Gottschalk *et al.* 2008). Some new anthophilic species were described recently for Neotropics (Vaz *et al.* 2014, 2018; Grimaldi 2016; Llangarí & Rafael 2017; Vilela & Prieto 2018), but none of them seem to be coespecific with the species of unknown identity reported in the present survey.

### The restrict flower-breeding species

The species included here are thought to breed exclusively or preferentially in flowers. They are never or rarely attracted to fruit-baited traps.

#### *Drosophila bromeliae* species group

This group showed to be the most speciose, abundant and widespread, found in all sites surveyed (Figure 1) and also the more taxonomically challenging. Altogether, eight species were found. All this species are cryptic in respect to external morphology, but can be discriminated by aedeagus morphology. In the time of sampling, just five species of *D. bromeliae* species group were described, and descriptions of terminalia were not available for the two species described by Sturtevant (1916, 1921). This impeded to recognise if any of the species found were coespecific to Sturtevant’s species and, so, to recognise which of them represented new species. Just *D. bromelioides* specimens could be clearly recognised based on the male terminalia of the lectotype depicted by Val (1982). After Grimaldi (2016) redescribed the Sturtevant’s specimens, another of the species found in this survey could be recognised as *D. bromeliae*. The identity of the six other species found here deserves a more careful study, since there are subtle to great differences in the morphology of aedeagus from the nine new species described by Grimaldi (2016) based mostly on Central American specimens. They are mentioned herein as types III, III’, IV, IV’, V and VI.

The species found in the present survey showed marked differences in host species (Table 1), from very generalist to ecologically specialised species. Individuals of the polyphagous species start to emerge from flowers around 10-13 days after the field collection. The *Solanum*-specialised species takes a little longer, around 13-16 days.

*Drosophila bromelioides* was the most conspicuous species and proved to be widely generalist, recorded in 42 host species, from 16 botanical families. Although rarely, this species is sometimes caught at low numbers even in banana-baited traps (De Toni *et al.* 2007; Gottschalk *et al.* 2007; Schmitz *et al.* 2007) and under constant attention can be reared in banana-agar medium in laboratory. However, flowers are clearly its preferred breeding site. It was found in souuthern localities, in the states of Rio Grande do Sul (new southernmost locality in Pelotas), Santa Catarina, Paraná and São Paulo. Other studies have recorded it also in Goiás, Minas Gerais and Bahia (Vilela & Mori 1999; Chaves & Tidon 2008; Roque & Tidon 2008). A morphologically similar species, *D. bromeliae*, was found in northern Brazilian localities (Pará, Pernambuco and Mato Grosso states) in the present study and reported to Caribbean, Central America and Ecuador by Grimaldi (2016). Although some earlier studies reported *D. bromeliae* or *D. bromelioides* in other localities, caution are needed, since the cryptic diversity revealed here and by Grimaldi (2016). As examples, there are records for *D. bromeliae* in Campos do Jordão (São Paulo) by Hsu (1949) and *D. bromelioides* in Salvador (Bahia) by Malogolowkin (1951). However, both precede the first description of aedeagus for *D. bromelioides* (Val 1982) and for *D. bromeliae* (Grimaldi 2016), so, it is advisable to treat such cases as doubtful. Based in the results of the present survey, *D. bromeliae* is probably also a polyphagous species, found in five host species, of five families, and also can be reared in banana-agar medium. As our sampling effort in northern Brazilian regions was lower, these numbers can not be directly compared to those of *D. bromelioides*. Future samplings in the region are necessary to clear the extent of the niche breadth of this species. It is probable that various other hosts can be found.

Based on morphology, types III and III’ are thought to be related species. The type III was found in the southern states of Rio Grande do Sul and Santa Catarina, while the type III’ was found in the northeastern state of Pernambuco. Although also polyphagous, being found in six host species from five families, the type III exploits a very less variety of host plants comparing to the sympatric *D. bromelioides*. In turn, the type III’ was found in only one host species, being more restrict that the sympatric species *D. bromeliae*. However, future sampling probably must find other host plants for it, since it was discovered breeding in *Thunbergia grandiflora*, that is a cultivated exotic plant. Presumably, it may occupy a similar niche to its southern sibling type III, *i. e.*, polyphagous but not so generalist as *D. bromelioides* and *D. bromeliae*. Both types III and III’ can be reared, although difficultly, from banana-agar medium in laboratory.

In southern localities, *D. bromelioides* and type III can occur is some hosts both sympatrically and synchronically, although seemingly with some different preferences. A great number of type III specimens were reared from *Brugmansia suaveolens*, where it generally outnumbered *D. bromelioides*. *Ipomoea alba* was an exception in its genus for apparently representing an important host for type III, whereas other species of the genus host almost exclusively *D. bromelioides*. The type III’ was also find co-occurring with the more generalist *D. bromeliae* in *Thunbergia grandiflora* flowers in Pernambuco. All four species showed to be able to breed in some exotic plants.

Contrasting to their more generalist siblings, four more specialised species complete the group, all of them using exclusively species of *Solanum* as host species. Despite some effort, no one of these species could be reared in artificial culture media in laboratory, even when some kind of *Solanum* flower juice was added to the medium. These species are being called here as types IV, IV’, V and VI.

The types IV and IV’ are morphologically similar. The type IV was found in the southern states of Rio Grande do Sul and Santa Catarina, while the type IV’ was found in Bahia, Minas Gerais, São Paulo, Paraná and Santa Catarina. Additionally, the type V was also found in the southern states of Rio Grande do Sul and Santa Catarina. This species share some hosts with type IV, although the two species seem to have some different host preferences. Both species breed in *Solanum paniculatum*, separately or together; in the other hand, type IV occurs solely in *S. mauritianum*, while type V is the only found in *S. sisymbriifolium* in these localities. Other *Solanum* species were less sampled, but both fly species shared *S. sanctaecatharinae*. The type V showed to have a niche slightly wider, being found in six species of *Solanum*, while the type IV was found in three species. Some specimens of the generalist *D. bromelioides* was reared from *Solanum* flowers, but in a few occasions and at low numbers, suggesting that these genus does not represent a main host for it. Finally, in a sample taken in the northern state of Pará, other species was found, the type VI. Its host plant, *S. stramoniifolium* does not occur in southern Brazil.

*Solanum* is a large and worldwide distributed genus, especially in tropical and subtropical regions (Mentz & Oliveira 2004). Species sampled in the present survey are native from Neotropics and are mostly ruderal plants, occurring in pioneer types of vegetation, in roadsides, field and forest borders. However, no equivalent *Solanum*-specialised drosophilid is known in other parts of the world. The only literature record known by us is on the occurrence of *Scaptodrosophila minimeta* in flowers of *S. torvum* and *S. mauritianum* in Australia (Bock & Parson 1981), but the ecology of this species is poorly known. It can be supposed that this species uses other natural resource, as both species of S*olanum* are native from Neotropics and introduced in Australia.

The records presented here suggest the existence of pairs of morphologically and ecologically similar species in different localities in Brazil, in each case with one of them found in northern localities and another in southern localities (the generalist *D. bromeliae* and *D. bromelioides*, the oligophagous types III’ and III, the *Solanum*-specialised types IV’ and IV, respectively with a northern and a southern distribution). Though, since our sampling in the huge Brazilian territory and plant biodiversity was not exhaustive and was biased to southern localities, this preliminary observation needs to be tested by future studies.

#### *Drosophila lutzii* species group (= subgenus *Phloridosa* of *Drosophila*)

This taxon comprises only Neotropical flower-breeding species, traditionally classified as the subgenus *Phloridosa* (Sturtevant 1942), but recently transferred to the subgenus *Drosophila* as *D. lutzii* species group (Yassin 2013). Two species were represented in our samples, *D. denieri* and *D. lutzii*. Their time of emergence of imagines was around 10-13 days after the flowers were collected. *Drosophila lutzii* are widespread in Southern United States, Mexico, Central America (Sturtevant 1921; Chassagnard & Tsacas 1992), and introduced in Hawaii (Montague & Kaneshiro 1982), but only scattered sampled in South America (Pilares & Suyo 1982, in Peru; Vilela 1984, in Argentina; Acurio & Rafael 2009, in Ecuador). It was recorded in Brazil for the first time by Schmitz and Hofmann (2005), in the southern states of Rio Grande do Sul and Santa Catarina, followed by Roque and Tidon (2008) in the states of Mato Grosso and Goiás, in central Brazil. In the present survey it is recorded for the first time in the states of São Paulo and Paraná. In turn, *D. denieri* is known only to South America, being referred for northern Argentina (Blanchard 1938), Uruguay (Goñi *et al.* 1998) and for the Brazilian states of Rio Grande do Sul, Santa Catarina (Schmitz & Hofmann 2005), Rio de Janeiro (Frota-Pessoa 1952) and Mato Grosso (Blauth & Gottschalk 2007). Both species are polyphagous. In the present survey, while *D. denieri* occurred in a wider variety of botanical families, including exotic elements, most specimens of *D. lutzii* occurred in *Ipomoea*. However, this species is also recorded for different botanical families (Chassagnard & Tsacas 1992; Roque & Tidon 2008). The two species co-occurred in *Brugmansia suaveolens*, but generally *D. denieri* was the most common species in this plant. In the other hand, *D. lutzii* was the most common species found in all species of *Ipomoea*, except *I. alba*. However, there are exceptions, as in Pelotas, where only *D. denieri* was found in *I. cairica* and *I. indica*. These two species, wherever found, almost always shared their host plants with species of the *Drosophila bromeliae* species group. Both species could be reared just very poorly in the banana-agar medium in laboratory, with just a few individuals for one generation or two, as previously reported for the group (Brncic, 1962b).

#### Zygothrica

It is reported that great numbers of adult flies of this genus can be found over bracket fungi in rainforests, but is questionable the kind of relation between flies and fungi. Undoubtedly, fungi actually represent breeding sites for some *Zygothrica* species (Grimaldi 1987; Roque *et al.* 2006; Gottschalk *et al.* 2009), but Grimaldi (1987) suggested that in many cases they are used only as rendezvous sites. Indeed, several records for breeding sites in the genus are flowers, although they are reported only for a few species (Grimaldi 1987; Fonseca *et al.* 2017). *Zygothrica dispar* is a good example of this behaviour: Burla (1954) stated that flies of this species were caught by sweeping a net over numerous fungi, but Frota-Pessoa (1952), Malogolowkin (1952) and Santos and Vilela (2005) reared the species from flowers. It was suggested before (Malogolowkin 1952; Santos & Vilela 2005) that this species is mainly a ground-feeder, using fallen flowers opportunistically. Contrarily, in our samples it was reared mainly in living flowers. Time for emergence was around 15-18 days. It demonstrated a preference for forested environments, being, in fact, the principal flower-breeding species in the area of Atlantic Forest of Morro da Lagoa da Conceição, Florianópolis. The other species, *Z.* sp.1, was reared only from *Solanum* flowers. Interestingly, this situation is analogous with that found in the *Drosophila bromeliae* species group, with *Solanum* again requiring a specialised performance.

### Species of uncertain biology

The nature of the interaction between the species included in the category “uncertain biology” and flowers of the host plants recorded in this survey is doubtful. These species are normally absent in baited trap-based surveys (De Toni *et al.* 2007; Schmitz *et al.* 2007; Döge *et al.* 2008; Garcia *et al.* 2012; and others). In the other hand, they were found only occasionally in our flower samples. If they are not specialised breeders in flowers, they may occupy still another unknown niche.

Both species of *Cladochaeta* Coquillett, 1900 found belong to the *C. nebulosa* species group. *Cladochaeta bupeo* was known previously only by a few individuals from Panama, collected during the decades of 1950s and 1960s (Grimaldi & Nguyen 1999). This finding represents, therefore, its first record in South America and its new southernmost locality (Porto Alegre, Rio Grande do Sul). The other species, *C.* sp.1, certainly is a related species, although undescribed. *Cladochaeta* remains a poorly-known genus in Brazil, as emphasised by Grimaldi & Nguyen (1999) in their extensive revision of the genus and by recent advances made by Pirani & Amorim (2016). The authors of both studies expect that many species of this genus remain to be recorded or described from Brazilian territory. The natural history of this genus remains still more obscure. Larvae of some species are reported to feed on immature stages of other arthropods (Grimaldi & Nguyen 1999; Carvalho-Filho *et al.* 2018). The possibility that the specimens from Porto Alegre just reached flowers to pupate after feeding on other resource can not be discarded. However, a few cases of association between *Cladochaeta* and flowers that may not involve Cercopidae hosts are reported: between *C. floridana* (Malloch, 1924) and *Bidens pilosa* (Asteraceae) in Florida, USA, between *C. psychotria* Grimaldi & Nguyen, 1999 and *Psychotria chiriquiensis* (Rubiaceae) in Costa Rica (Grimaldi & Nguyen 1999), and between *C.* aff. *paradoxa* and *Cestrum sendtnerianum* (Solanaceae) in São Paulo, Brazil (Santos & Vilela 2005). The two species found in Porto Alegre were represented only by two and three individuals, respectively, which emerged 12-13 days after flowers were collected, and were not found again in subsequent samples from the same plants.

The genus *Rhinoleucophenga* Hendel, 1917 was represented by one species found in Florianópolis, still undescribed in the occasion. Based on the seven specimens caught, it was subsequently described as *R. joaquina* (Schmitz *et al*. 2009). Its life cycle is considerably longer than other drosophilids reported here, as imagines took 20-21 days to emerge after the field collection of the flowers. Similarly to *Cladochaeta,* the natural history of *Rhinoleucophenga* is also poorly understood, but there are reports of larvae showing predacious behaviour on other insects (Costa Lima 1935; Vidal *et al.* 2018). So, the possibility that such specimens were attacking another insects near or inside the flowers can not be discarded. A subsequent attempt to collect it from the same plant and locality was unsuccessful, but a few specimens were sampled in banana-baited traps in Bahia and Rio Grande do Sul (Poppe *et al.* 2015).

Although being one of the larger genera of Drosophilidae, *Scaptomyza* is virtually ignored in Brazil, with just three described species recorded (Gottschalk *et al.* 2008). Three species were found in Porto Alegre and probably represent undescribed species. The time for emergence of imagines after collection ranged from 11-14 days for all species. *Scaptomyza* sp.1 (36 specimens) and *S.* sp.3 (15 specimens) belong to subgenus *Mesoscaptomyza* Hackman, 1959, while *S.* sp.2 (26 specimens) belongs to the genus *Parascaptomyza* Duda, 1924. Specimens of this genus were mainly found during late winter of 2006, in the flowering period of some Bignoniaceae, but were not found thereafter. Although the biology of the genus is not wholly understood, some species of *Scaptomyza* are known to be important flower breeders, especially in Hawaii, but can also feed on sap flood or leaves (Montague & Kaneshiro 1982; Magnacca *et al.* 2008). In Brazil, Santos and Vilela (2005) reported one individual of *Scaptomyza* sp. from flowers of *Cestrum sendtnerianum* (Solanaceae), in São Paulo.

### Opportunist species

Several species found in the present survey are not supposed to present ecological niches strictly linked to flowers. Most of them are common members of the Neotropical fruit assemblages of drosophilids (De Toni *et al.* 2001; Silva *et al.* 2005) and are usually attracted to fruit baited traps (Tidon 2006; De Toni *et al.* 2007; Gottschalk *et al.* 2007; Garcia *et al.* 2012), as *D. cardinoides, D. griseolineata, D. mediostriata, D. paulistorum, D. melanogaster, D. simulans* and *Zaprionus indianus*, the last three being exotic species. The only specimen classified as *D. tripunctata* gr. sp.1 was a female from *D. tripunctata* species group, not identified at species level, since there is a lot of cryptic species in the group, discriminated only by the males, but probably also fits this situation. Except for *D. melanogaster* and *D. paulistorum*, all of these species were reared exclusively from fallen flowers. Fallen flowers constitute a substrate very distinct from living flowers, requiring a less specialised performance (Brncic 1983), giving opportunities for generalist species, what Pipkin *et al.* (1966) called “ground-feeders”. Furthermore, the specimens of *D. paulistorum* reared from living flowers of *Allamanda polyantha* presented a strikingly reduced body size, suggesting that they passed the preadult stage at suboptimal conditions. Interestingly, specimens of *D. melanogaster* reared from living flowers of *Thunbergia grandiflora* (from Recife) presented a *brown*-like phenotype. Other evidence that suggests that these species use flowers only as a secondary or incidental host is the fact that they were not found repeatedly in such hosts. *Drosophila cardinoides* was abundantly reared (n>100) in Porto Alegre from fallen flowers of *Ceiba speciosa* in late Summer of 2007, which demonstrates its ability to exploit such resource, but no other individual were found in subsequent samples of the same tree, even in the same site, indicating that this association is not constant. Other species were found in lower numbers (normally n<10). The only species found repeatedly in a host plant was *D. mediostriata* in fallen flowers of *Hibiscus pernambucensis*. This species is, nevertheless, also reared from fruits (Heed 1957; De Toni *et al.* 2001), although its presence in flowers was already reported (Dobzhansky & Pavan 1950; Pikin *et al.* 1966). Unusually, Schmitz *et al.* (2007) found this species abundantly in banana baited traps at mangrove forests of Santa Catarina Island, southern Brazil, contrasting to very lower abundances in similar surveys in other environments of the same region (De Toni *et al.* 2007; Gottschalk *et al.* 2007; Bizzo *et al.* 2010). *Hibiscus pernambucensis* is one of the main floristic components of peripheral areas of the mangroves in southern Brazil. However, *D. mediostriata* oviposits exclusively in fallen flowers, since only *D. bromelioides* was reared from samples of living flowers, showing a clear niche separation of the two species in the flowers of this plant.

The remaining two species are less common in diversity surveys, but still there are evidences that they are not specialised in flowers. *Drosophila mediovitatta* was already reported from flowers before (Goñi *et al.* 1998), but also from rotten cladodes (Goñi *et al.* 1998) and from fungi (Gottschalk *et al.* 2009), this last record in the same locality of the present study (Porto Alegre). Finally, *D. sticta* was also recorded from flowers before (Pipkin *et al.* 1966; Santos & Vilela 2005), but also in banana-baited traps (Medeiros & Klaczko 2004; Garcia *et al.* 2012).

### Future directions

Our sampling was far to be exhaustive, and biased to southern localities. The huge territorial extension of Brazil and the extremely high spatial heterogeneity and diversity of flora make a complete assessment of the flower-breeding diversity of drosophilids a challenging task. So, there is a high demand for new efforts in searching for flower-breeding drosophilids in Brazil and Neotropical Region as a whole, widening both the range of geographical localities and the diversity of flora surveyed as potential hosts. Several other Drosophilidae taxa not sampled in the present survey are known to be flower-breeding in Neotropical region, as the *D. onychophora* and *D. xanthopallescens* species groups, additional species of the *D. bromeliae* and *D. lutzii* species groups and some species associated to Araceae. Even the *Drosophila flavopilosa* species group, a comparatively better studied taxon and not targeted here, still requires better understanding. Surveys on Drosophilidae diversity are still highly needed, and alternative methods of sampling must be considered besides the traditional banana-baited traps and fallen fruits. Collecting drosophilids from flowers is a relatively simple and cheap method, so it is not difficult to be included in diversity inventories. In the other hand, as shown here, the taxonomic impediment is still very high and probably the most prohibitive challenge for new researches. It is quite certain that many Neotropical flower-breeding species still wait for being discovered and integrative taxonomy that links morphological and molecular characters will be probably needed to guide the understanding of species delimitations.

Identifying properly the taxa linked to flowers in a satisfactory biogeographical and ecological framework is a prerequisite to elucidate episodes of host shifts and the repeated evolution of flower-breeding habits within family Drosophilidae. Understanding the natural history of these insects may provide important highlights on the evolution of specialisation to host plants. The nature of the insect/plant interaction remains to be unveiled. Reliable information on what exactly the larvae and adults feed would be welcome. Literature reports on larvae or adults feeding on plant tissues or fluids, pollen or yeasts are more conjectural that empirical. Other issues to be addressed are the role of potentially toxic plant secondary compounds as selective factors for the adaptive process of colonizing a new host species, besides the chemosensory mechanisms related to the insect recognition of hosts, especially those with narrower niches. A multi-disciplinary approach would be essential for giving a light to these questions.

The occurrence of species with a broad range of generalist to specialist niches in the *Drosophila bromeliae* species group, as well as preliminary evidences of differences in the geographical distribution of the species, provides an interesting opportunity to study events of speciation, involving potentially biogeographical or ecological factors.

Studies on tropical assemblages of drosophilids normally are challenged by the huge diversity and complexity of such associations. The study of flower-breeding drosophilids is an opportunity to study the species coexistence through a relatively simpler model. Other ecological, genetic and evolutionary mechanisms could be better understood. As Brncic (1983) stated many years ago, the study of species with restricted niches like the flower-breeding forms is advantageous because it is possible to relate their genetic structure with the ecology under simpler variables, so allowing a better understanding of the evolutionary process of the members of the family Drosophilidae as a whole.

## ACKNOWLEDGMENTS

People from various institutions provided HJS with valuable help in collection trips, sending material or making available their labs: Monica L. Blauth (Universidade Federal de Pelotas), Marco S. Gottschalk and Marlúcia B. Martins (Museu Paraense Emílio Goeldi), Cláudia Rohde, Ana Cristina L. Garcia (Universidade Federal de Pernambuco, Centro Acadêmico de Vitória), José Ferreira dos Santos, Tania T. Rieger, Evilis S. Monte, Luciana F. de Souto (Universidade Federal de Pernambuco, Centro de Ciências Biológicas) and Paulo R. P. Hofmann (Universidade Federal de Santa Catarina). The following people kindly identified some plant species: Lilian A. Mentz, João André Jarenkow (Universidade Federal do Rio Grande do Sul), Luiz Carlos B. Lobato, Francismeire Bonadeu da Silva and Ana Kelly Kock (Museu Paraense Emílio Goeldi). Sincere thanks are due to all of them and to the team of Laboratório de *Drosophila* at Universidade Federal do Rio Grande do Sul. HJS is in dept to David Grimaldi for his review on *Drosophila bromeliae* species group, without which the present survey could not go further properly. This research was supported by grants and fellowships of Conselho Nacional de Desenvolvimento Científico e Tecnológico (CNPq).

## Supporting Information

Additional Supporting Information may be found in the online version of this article: Appendix S1 Plant species with no emergence of drosophilids.

### Supporting Information

**Appendix S1:** List of plant species with no emergence of drosophilid specimens in the present survey.

|  |  |
| --- | --- |
| **Acanthaceae** | *Justicia floribunda* |
|  | *Sanchezia oblonga* |
|  | *Odontonema strictum* |
|  | sp.1 |
| **Amaryllidaceae** | *Agapanthus africanus* |
| **Asteraceae** | *Mikania cordifolia* |
|  | *Senecio brasiliensis* |
| **Apocynaceae** | *Allamanda blanchetii* |
|  | *Nerium oleander* |
|  | *Tabernaemontana catharinensis* |
|  | *Thevetia peruviana* |
| **Araceae** | sp.1 |
| **Balsaminaceae** | *Impatiens walleriana* |
| **Begoniaceae** | *Begonia coccinea* |
| **Bignoniaceae** | *Adenocalymma schomburgkii* |
|  | *Podranea ricasoliana* |
| **Cactaceae** | *Opuntia cochenillifera* |
| **Cannaceae** | *Canna indica* |
|  | *Canna paniculata* |
| **Cleomaceae** | *Tarenaya hassleriana* |
| **Convolvulaceae** | *Ipomoea imperati* |
| **Commelinaceae** | *Tradescantia* sp. |
| **Cucurbitaceae** | *Momordica charantia* |
| **Ericaceae** | *Gaylussacia brasiliensis* |
|  | *Rhododendron simsii* |
| **Escalloniaceae** | *Escallonia bifida* |
| **Euphorbiaceae** | *Euphorbia pulcherrima* |
| **Fabaceae** | *Bauhinia forficata* |
|  | *Delonix regia* |
|  | *Lupinus* sp. |
|  | *Macroptilium lathyroides* |
|  | *Peltophorum dubium* |
|  | *Schizolobium parahyba* |
| **Geraniaceae** | *Pelargonium x hortorum* |
| **Heliconiaceae** | *Heliconia farinosa* |
| **Iridaceae** | *Gladiolus x hortulanum* |
| **Lamiaceae** | *Salvia coccinea* |
| **Liliaceae** | *Lilium regale*<br>*Lilium* sp.1 |
| **Lythraceae** | *Lagerstroemia indica*<br>*Punica granatum* |
| **Magnoliaceae** | *Magnolia liliflora* |
| **Malvaceae** | *Abelmoschus esculentus*<br>*Brachychiton populneus*<br>*Hibiscus schizopetalum*<br>*Luehea divaricata* |
| **Melastomataceae** | *Tibouchina urvilleana* |
| **Musaceae** | *Musa ornata* |
| **Nymphaeaceae** | *Nymphaea ampla* |
| **Oleaceae** | *Jasminum mesnyi* |
| **Onagraceae** | *Ludwigia* sp. |
| **Orchidaceae** | *Epidendrum fulgens*<br>sp.1 |
| **Oxalidaceae** | *Oxalis corniculata*<br>*Oxalis latifolia* |
| **Piperaceae** | *Piper* sp. |
| **Plantaginaceae** | *Russelia equisetiformis*<br>sp.1 |
| **Proteaceae** | *Grevillea banksii* |
| **Rosaceae** | *Rosa spinosissima* |
| **Solanaceae** | *Datura stramonium*<br>*Petunia integrifolia*<br>*Petunia* sp.<br>*Solanum pycnanthemum*<br>*Streptosolen jamesonii* |
| **Urticaceae** | *Cecropia pachystachya* |
| **Verbenaceae** | *Lantana camara*<br>*Petrea subserrata* |
| **Zingiberaceae** | *Renealmia* cf. *petasites* |

## REFERENCES

Acurio A, Rafael V (2009) Diversity and geographical distribution of *Drosophila* (Diptera, Drosophilidae) in Ecuador. Drosophila Information Service 92, 20–25.

APG IV (2016) An update of the Angiosperm Phylogeny Group classification for the orders and families of flowering plants: APG IV. Botanical Journal of the Linnean Society 181, 1–20.

Bächli G, Vilela CR, Escher SA, Saura A (2004) The Drosophilidae (Diptera) of Fennoscandia and Denmark. Fauna Entomologica Scandinavica 39, 1–362.

Bizzo LEM, Gottschalk MS, De Toni DC, Hofmann PRP (2010) Seasonal dynamics of a drosophilid assemblage and its potential as bioindicators in open environments. Iheringia, Série Zoologia 100, 185–191.

Blanchard EE (1938) Descripciones y anotaciones de Dipteros argentinos. Anales de la Sociedad Científica Argentina 126, 345–386.

Blauth ML, Gottschalk MS (2007) A novel record of Drosophilidae species in the Cerrado biome of the state of Mato Grosso, west-central Brazil. Drosophila Information Service 90, 90–96.

Bock IR, Parsons PA (1981) Species of Australia and New Zealand. In: Ashburner M, Carson HL, Thompson, Jr. JN (eds) The Genetics and Biology of Drosophila. Vol. 3a, pp. 291–308. Academic Press, London.

Brncic D (1962a) Chromosomal structure of populations of *Drosophila flavopilosa* studied in larvae collected in their natural breeding sites. Chromosoma 13, 183–195.

Brncic D (1962b) New Chilean species of the genus *Drosophila*. Biologica 33, 3–6.

Brncic D (1966) Ecological and cytogenetical studies of *Drosophila flavopilosa*, a Neotropical species living in *Cestrum flowers*. Evolution 20, 16–29.

Brncic D (1978) A note on the *flavopilosa* group of species of *Drosophila* in Rio Grande do Sul, Brazil, with description of two new species (Diptera, Drosophilidae). Revista Brasileira de Biologia 38, 647–651.

Brncic D. (1983) Ecology of flower-breeding *Drosophila*. In: Ashburner M, Carson HL, Thompson, Jr. (eds) The genetics and biology of Drosophila, Vol. 3d, pp 333–382. Academic Press, London.

Brown BV (2005) Malaise trap catches and the crisis in Neotropical dipterology. American Entomologist 51, 180–183.

Burla H (1954) Study on the polymorphism in *Zygothrica dispar* and *Z. prodispar*, and description of *Z. laticeps* sp. nov. (Drosophilidae, Diptera). Arquivos do Museu Paranaense 10, 231–252.

Carson HL, Okada T (1980) Drosophilidae associated with flowers in Papua New Guinea. I. Colocasia esculenta. Kontyu 48, 15–29.

Carvalho-Filho FDS, Pirani G, Kloss TG (2018) A new species and notes on unusual natural history of *Cladochaeta* Coquillett, 1900 (Diptera: Drosophilidae). Zootaxa 4410, 483–496.

Chassagnard MT, Tsacas L (1992) *Drosophila* (*Phloridosa*) *lutzii* Sturtevant (Diptera: Drosophilidae), espécie antófila de México. Folia Entomológica Mexicana 85, 95–105.

Chaves NB, Tidon R (2008) Biogeographical aspects of drosophilids (Diptera, Drosophilidae) of the Brazilian savanna. Revista Brasileira de Entomologia 52, 340–348.

Cook RM, Parsons PA, Bock IR (1977) Australian endemic *Drosophila*. II. A new *Hibiscus*-breeding species with its description. Australian Journal of Zoology 25, 755–763.

Costa Lima, A (1935) Um Drosophilideo predador de Coccideos. Chácaras e Quintaes 52, 61–63.

De Toni DC, Gottschalk MS, Cordeiro J, Hofmann PRP, Valente VLS (2007) Study of the Drosophilidae (Diptera) Communities on Atlantic Forest Islands of Santa Catarina State, Brazil. Neotropical Entomology 36, 356–375.

De Toni DC, Hofmann PRP, Valente VLS (2001) First record of *Zaprionus indianus* (Diptera, Drosophilidae) in the State of Santa Catarina, Brazil. Biotemas 14, 71–85.

Dobzhansky T, Pavan C (1943) Studies on Brazilian species of *Drosophila*. Boletim da Faculdade de Filosofia, Ciências e Letras da Universidade de São Paulo 36, 7–72.

Dobzhansky T, Pavan C (1950) Local and seasonal variations in relative frequencies of species of *Drosophila* in Brazil. Journal of Animal Ecology 19, 1–14.

Döge JS, Valente VLS, Hofmann PRP (2008) Drosophilids (Diptera) from an Atlantic Forest area in Santa Catarina, Southern Brazil. Revista Brasileira de Entomologia 52, 615–624.

Duda O (1925) Die costaricanischen Drosophiliden des Ungarischen National-Museums zu Budapest. Annales Musei Nationalis Hungarici 22, 149–229.

Duda O (1927) Die sudamerikanischen Drosophiliden (Dipteren) unter Berucksichtigung auch der anderen neotropischen sowie der nearktischen Arten. Archiv fur Naturgeschichte 91, 1–228.

Endara L, Grimaldi DA, Roy BA (2010) Lord of the flies: pollination on *Dracula* orchids. Lankesteriana 10, 1–11.

Fartyal RS, Gao J-J, Toda MJ, Hu Y-G, Takenaka Takano K, Suwito A, Katoh T, Takigahira T, Yin J-T (2013) *Colocasiomyia* (Diptera: Drosophilidae) revised phylogenetically, with a new species group having peculiar lifecycles on monsteroid (Araceae) host plants. Systematic Entomology 38, 763–782.

Fonseca PM, Loreto ELS, Gottschalk MS, Robe LJ (2017) Cryptic diversity and speciation in the *Zygothrica* genus group (Diptera, Drosophilidae): the case of *Z. vittimaculosa* Wiedemann. Insect Systematics & Evolution 48, 285–313.

Frota-Pessoa O (1952) Flower-feeding Drosophilidae. Drosophila Information Service 26, 101–102.

Garcia CF, Hochmüller CJC, Valente VLS, Schmitz HJ (2012) Drosophilid assemblages at different urbanization levels in the city of Porto Alegre, state of Rio Grande do Sul, southern Brazil. Neotropical Entomology 41, 32–41.

Goñi B, Martinez ME, Valente VLS, Vilela CR (1998) Preliminary data on the *Drosophila* species (Diptera, Drosophilidae) from Uruguay. Revista Brasileira de Entomologia 42, 131–140.

Gottschalk MS, Bizzo LEM, Döge JS, Profes MS, Hofmann PRP, Valente (2009) Drosophilidae (Diptera) associated to fungi: differential use of resources in anthropic and Atlantic Rain Fores areas. Iheringia, Série Zoologia 99, 442–448.

Gottschalk MS, De Toni DC, Valente VLS, Hofmann PRP (2007) Changes in Brazilian Drosophilidae (Diptera) Assemblages Across an Urbanization Gradient. Neotropical Entomology 36, 848–862.

Gottschalk MS, Hofmann PRP, Valente VLS (2008) Diptera, Drosophilidae: historical occurrence in Brazil. Check List 4, 485–518.

Grimaldi DA, Nguyen T (1999) Monograph on the spittlebug flies, genus *Cladochaeta* (Diptera: Drosophilidae: Cladochaetini). Bulletin of the American Museum of Natural History 241, 1–326.

Grimaldi DA (1987) Phylogenetics and taxonomy of *Zygothrica* (Diptera: Drosophilidae). Bulletin of the American Museum of Natural History 186, 103–268.

Grimaldi DA (2016) Revision of the *Drosophila bromeliae* species group (Diptera: Drosophilidae): Central American, Caribbean, and Andean Species. American Museum Novitates 3859, 1–55.

Heed WB (1957) Ecological and distributional notes on the Drosophilidae (Diptera) of El Salvador. The University of Texas Publication 5721, 62–78.

Heed WB, Carson HL, Carson MS (1960) A list of flowers utilized by drosophilids in the Bogota region of Colombia. Drosophila Information Service 34, 84–85.

Hsu TC (1949) The external genital apparatus of male Drosophilidae in relation to systematics. The University of Texas Publication 4920, 80–142.

Hunter AS (1979) New anthophilic *Drosophila* of Colombia. Annals of the Entomological Society of America 72, 372–383.

Llangarí L, Rafael V (2017) A new species of *Drosophila* (Diptera: Drosophilidae) from the inflorescences of *Xanthosoma sagittifolium* (Araceae). Revista Ecuatoriana de Medicina y Ciencias Biologicas 38, 55–62.

Magnacca KN, Foote D, O’Grady PM (2008) A review of the endemic Hawaiian Drosophilidae and their host plants. Zootaxa 1728, 1–58.

Malogolowkin C (1951) Drosophilideos colhidos na Bahia, com descrição de uma espécie nova (Diptera). Revista Brasileira de Biologia 11, 431–434.

Malogolowkin C (1952) Notas sobre *Zygothrica dispar* (Diptera, Drosophilidae). Revista Brasileira de Biologia 12, 455–457.

May RM (2010) Ecological science and tomorrow’s world. Philosophical Transactions of the Royal Society, B 365, 41–47.

McEvey SF, Barker JSF (2001) *Scaptodrosophila aclinata*: a new *Hibiscus* flower-breeding species related to *S. hibisci* (Diptera: Drosophilidae). Records of the Australian Museum 53, 255–262.

Medeiros HF, Klaczko LB (2004) How many species of *Drosophila* (Diptera, Drosophilidae) remain to be described in the forests of São Paulo, Brazil? Species lists of three forest remnants. Biota Neotropica 4, 1–12.

Mentz LA, Oliveira PL (2004) *Solanum* (Solanaceae) na Região Sul do Brasil. Pesquisas. Botânica 54, 1–327.

Montague JR, Kaneshiro KY (1982) Flower-breeding species of Hawaiian Drosophilids in an early stage of sympatry. Pacific Insects 24, 209–213.

Myers N, Mittermeier RA, Mittermeier CG, Fonseca GAB, Kent J (2000) Biodiversity hotspots for conservation priorities. Nature 403, 853–858.

Pavan C, Cunha AB (1947). Espécies brasileiras de Drosophila. Boletim da Faculdade de Filosofia, Ciências e Letras da Universidade de São Paulo 86, 20–64.

Pilares GLV, Suyo MP (1982) Distribution of different species of *Drosophila* from Peru (South America). Drosophila Information Service 58, 122–124.

Pipkin SB (1964) New flower breeding species of *Drosophila*. Proceedings of the Entomological Society Of Washington 66, 217–245.

Pipkin SB, Rodríguez RL, León J (1966) Plant host specificity among flower-feeding neotropical *Drosophila* (Diptera: Drosophilidae). The American Naturalist 100, 135–155.

Pirani G, Amorim DS (2016) Going beyond the tip of the Drosophilidae iceberg: new *Cladochaeta* Coquillett, 1900 (Diptera: Drosophilidae) from Brazil. Zootaxa 4139, 301–44.

Poppe JL, Schmitz HJ, Valente VLS (2015) The New World genus *Rhinoleucophenga* (Diptera: Drosophilidae): new species and notes on occurrence records. Zootaxa 3955, 349–370.

Powell JR (1997) Progress and Prospects in Evolutionary Biology. The Drosophila Model. Oxford University Press, Oxford.

Rafael JA, Aguiar AP, Amorim DS (2009) Knowledge of insect diversity in Brazil: challenges and advances. Neotropical Entomology 38, 565–570.

Robe LJ, De Ré FC, Ludwig A, Loreto ELS (2013) The *Drosophila flavopilosa* species group (Diptera, Drosophilidae): an array of exciting questions. Fly 7, 59–69.

Roque F, Figueiredo R, Tidon R (2006) Nine new records of drosophilids in the Brazilian savanna. Drosophila Information Service 89, 14–17.

Roque F, Tidon R (2008) Eight new records of drosophilids (Insecta; Diptera) in the Brazilian savanna. Drosophila Information Service 91, 94–98.

Santos RCO, Vilela CR (2005) Breeding sites of Neotropical Drosophilidae (Diptera). IV. Living and fallen flowers of *Sessea brasiliensis* and *Cestrum* spp. (Solanaceae). Revista Brasileira de Entomologia 49, 544–551.

Schmitz HJ, Hofmann PRP (2005) First record of subgenus *Phloridosa* of *Drosophila* in southern Brazil, with notes on breeding sites. Drosophila Information Service 88, 97–101.

Schmitz HJ, Valente VLS, Hofmann PRP (2007) Taxonomic survey of Drosophilidae (Diptera) from mangrove forests of Santa Catarina Island, southern Brazil. Neotropical Entomology 36, 53–64.

Schmitz HJ, Gottschalk MS, Valente VLS (2009) *Rhinoleucophenga joaquina* sp. nov. (Diptera: Drosophilidae) from the Neotropical Region. Neotropical Entomology 38, 786–790.

Silva AAR, Martins MB (2004) A new anthophilic species of *Drosophila* Fallén belonging to the *bromeliae* group of species (Diptera, Drosophilidae). Revista Brasileira de Zoologia 21, 435–437.

Silva NM, Fantinel CC, Valente VLS, Valiati VH (2005) Ecology of colonizing populations of the figfly *Zaprionus indianus* (Diptera, Drosophilidae) in Porto Alegre, southern Brazil. Iheringia, Série Zoologia 95, 233–240.

Souza VC, Lorenzi H (2005) Botânica Sistemética – guia ilustrado para identificação das famílias de angiospermas da flora brasileira, baseado em APG II. Plantarum, Nova Odessa.

Starmer WT, Bowles JM (1994) The saptial distribution of endemic and introduced flower-breeding species of *Drosophila* (Diptera: Drosophilidae) during their early history of encounter on the islands of Hawaii. Pan-Pacific Entomologist 70, 230–239.

Sturtevant AH (1916) Notes on North American Drosophilidae with descriptions of twenty-three new species. Annals of the Entomological Society of America 9, 323–343.

Sturtevant AH (1921) The North American species of *Drosophila*. Carnegie Institute of Washington Publication 301, 1–150.

Sturtevant AH (1942) The classification of the genus *Drosophila*, with descriptions of nine new species. The University of Texas Publication 4213, 5–51.

Sultana F, Hu YG, Toda MJ, Takenaka K, Yafuso M (2006) Phylogeny and classification of *Colocasiomyia* (Diptera, Drosophilidae), and its evolution of pollination mutualism with aroid plants. Systematic Entomology 31, 684–702.

Sultana F, Kimura MT, Toda MJ (1999) Anthophilic *Drosophila* of the *elegans* species-subgroup from Indonesia, with description of a new species (Diptera: Drosophilidae). Entomological Science 2, 121–126.

Suwito F, Kimura MT, Hattori K, Kimura MT (2002) Environmental adaptations of two flower breeding species of *Drosophila* from Java, Indonesia. Entomological Science 5, 399–406.

Tidon R (2006) Relationships between drosophilids (Diptera, Drosophilidae) and the environment in two contrasting tropical vegetations. Biological Journal of the Linnean Society 87, 233–247.

Tsacas L, Chassagnard MT, David JR (1988) Un nouveau groupe d’espèces afrotropicales anthophiles dans le sous-genre *Scaptodrosophila* du genre *Drosophila* (Diptera, Drosophilidae). Annales de la Societé Entomologique de France 24, 181–202.

Tsacas L, Chassagnard M-T (1992) Les relations Araceae-Drosophilidae. Drosophila aracea une espèce anthophile associée à l’Aracée *Xanthosoma robustum* au Mexique (Diptera: Drosophilidae). Annales de la Société entomologique de France 28, 421–439.

Val FC (1982) The male genitalia of some Neotropical Drosophila: notes and illustrations. Papéis Avulsos de Zoologia 34, 309–347.

Vaz SC, Vilela CR, Krsticevic FJ, Carvalho AB (2014) Developmental sites of Neotropical Drosophilidae (Diptera): V. Inflorescences of *Calathea cylindrica* and *Calathea monophylla* (Zingiberales: Marantaceae). Annals of the Entomological Society of America 107, 607–620.

Vaz SC, Vilela CR, Carvalho AB (2018) Two new species of *Drosophila* (Diptera, Drosophilidae) associated with inflorescences of *Goeppertia monophylla* (Marantaceae) in the city of São Paulo, state of São Paulo, Brazil. Revista Brasileira de Entomologia 62, 159–168.

Vidal MC, Sendoya SF, Yamaguchi LF, Kato MJ, Oliveira RS, Oliveira PS (2018) Natural history of a sit-and-wait dipteran predator that uses extrafloral nectar as prey attractant. Environmental Entomology 47, 1165–1172.

Vilela CR (1984) Notes on the holotypes of four Neotropical species of the genus *Drosophila* (Diptera, Drosophilidae) described by A. H. Sturtevant. Revista Brasileira de Entomologia 28, 245–256.

Vilela CR (1986) The type-series of *Drosophila denieri* Blanchard (Diptera, Drosophilidae). Revista Brasileira de Entomologia 30, 223–226.

Vilela CR, Mori L (1999) The genus *Drosophila* (Diptera, Drosophilidae) in the Serra do Cipó: further notes. Revista Brasileira de Entomologia 43, 319–128.

Vilela CR, Prieto D (2018) A new Costa Rican species of *Drosophila* visiting inflorescences of the hemi-epiphytic climber *Monstera lentii* (Araceae). Revista Brasileira de Entomologia 62, 225–231.

Yafuso M, Toda MJ, Sembel DT (2008) *Arengomyia*, new genus for the *Colocasiomyia arenga* species group (Diptera: Drosophilidae), with description of a new species. Entomological Science 11, 391–400.

Yassin A (2013) Phylogenetic classification of the Drosophilidae Rondani (Diptera): the role of morphology in the postgenomic era. Systematic Entomology 38, 349–364.

